# Precision pharmacological reversal of genotype-specific diet-induced metabolic syndrome in mice informed by transcriptional regulation

**DOI:** 10.1101/2023.04.25.538156

**Authors:** Phillip Wulfridge, Adam Davidovich, Anna C. Salvador, Gabrielle C. Manno, Rakel Tryggvadottir, Adrian Idrizi, M. Nazmul Huda, Brian J. Bennett, L. Garry Adams, Kasper D. Hansen, David W. Threadgill, Andrew P. Feinberg

**Affiliations:** Center for Epigenetics, Johns Hopkins School of Medicine, 855 N. Wolfe St, Baltimore, MD 21205, USA; Department of Biomedical Engineering, Johns Hopkins University, Baltimore, MD, USA; Department of Molecular and Cellular Medicine, Texas A&M Health Science Center, College Station, TX, USA; Department of Nutrition, Texas A&M University, College Station, TX, USA; Department of Nutrition, University of California, Davis, CA, US; Obesity and Metabolism Research Unit, USDA, ARS, Western Human Nutrition Research Center, Davis, CA, USA; Department of Veterinary Pathobiology, Texas A&M University, College Station, TX, USA; Department of Biostatistics, Johns Hopkins Bloomberg School of Public Health, 615 N. Wolfe St, Baltimore, MD 21205, USA; McKusick-Nathans Institute of Genetic Medicine, Johns Hopkins School of Medicine, 733 N. Broadway, Baltimore, MD 21205, USA; Department of Biochemistry & Biophysics, Texas A&M University, College Station, TX, USA; Department of Medicine, Johns Hopkins School of Medicine, 855 N. Wolfe St, Baltimore, MD 21205, USA; Department of Mental Health, Johns Hopkins Bloomberg School of Public Health, 624 N. Broadway, MD 21205, USA

## Abstract

Diet-related metabolic syndrome is the largest contributor to adverse health in the United States. However, the study of gene-environment interactions and their epigenomic and transcriptomic integration is complicated by the lack of environmental and genetic control in humans that is possible in mouse models. Here we exposed three mouse strains, C57BL/6J (BL6), A/J, and NOD/ShiLtJ (NOD), to a high-fat high-carbohydrate diet, leading to varying degrees of metabolic syndrome. We then performed transcriptomic and genomic DNA methylation analyses and found overlapping but also highly divergent changes in gene expression and methylation upstream of the discordant metabolic phenotypes. Strain-specific pathway analysis of dietary effects reveals a dysregulation of cholesterol biosynthesis common to all three strains but distinct regulatory networks driving this dysregulation. This suggests a strategy for strain-specific targeted pharmacologic intervention of these upstream regulators informed by transcriptional regulation. As a pilot study, we administered the drug GW4064 to target one of these genotype-dependent networks, the Farnesoid X receptor pathway, and found that GW4064 exerts genotype-specific protection against dietary effects in BL6, as predicted by our transcriptomic analysis, as well as increased inflammatory-related gene expression changes in NOD. This pilot study demonstrates the potential efficacy of precision therapeutics for genotype-informed dietary metabolic intervention, and a mouse platform for guiding this approach.

## Main

The extension of personalized medicine, the configuration of therapeutic regimens on an individual basis, will be critical for addressing public health issues, especially those related to environmental exposure. The importance of genotype and the epigenome in mediating phenotypic responses to environmental factors, and the overwhelming diversity of individual responses compared to population-level measurements, has become increasingly clear in recent years [1-4]. One such example is the known role of genotype in modulating how diet contributes to obesity and metabolic syndrome [5, 6]. Such "gene-by-diet", or GxD, interactions have been shown to explain, in great part, why a dietary recommendation that is beneficial for one demographic may be ineffective or even deleterious in another [7-9]. Personalized nutritional guidelines are therefore one promising solution to addressing obesity, though constructing such guidelines will require a continued effort towards establishing an in-depth understanding of GxD effects as well as their epigenetic and transcriptomic bases.

Despite the clear importance of identifying GxD interactions governing health effects in humans, there are many limitations to studying GxD in human cohorts [10]. For example, dietary backgrounds and habits of individuals are highly diverse and variable over a lifetime, translating to a lack of control over prior exposures and the need for extremely large cohort sizes in GxD studies to compensate for substantial confounding factors and noise. Conversely, controlled experiments on volunteers involving strict dietary regimens are highly likely to suffer from compliance issues as well as noise from variation in other environmental factors. Moreover, many GxD effects may be specifically mediated through the epigenome of metabolically-relevant tissues such as the liver and pancreas [11, 12], but obtaining samples of these tissues from human patients is often difficult or infeasible. Because human studies face these numerous challenges, animal models represent a powerful alternative for studying GxD with the crucial advantage of allowing extensive control and reproducibility of the experimental design. We have previously shown both that epigenetic analysis of mice reveals patterns of genetic susceptibility that are conserved in humans [13] and that genetically diverse mice have highly disparate phenotypic responses to diet [7].

The principles of GxD interactions, in which genotype is a key determinant of response to a particular diet or nutrient, also extend to other environmental exposures. Notably, they may even apply to the efficacy of therapeutic interventions such as the administration of drugs to alleviate metabolic disease. Currently, mouse studies that test the efficacy and safety of therapeutic regimens generally use only one laboratory strain such as C57BL/6J (BL6), which restricts their applicability to genetically diverse human patient populations. That is, a drug that happens to benefit the one specific mouse strain tested may nevertheless appear to fail efficacy tests in a larger clinical trial if other genetic backgrounds are insensitive to the drug; conversely, "rare" adverse effects that are actually genotype-dependent may be undetectable in one mouse strain [14]. Therefore, the use of multiple genetically distinct mouse strains in drug trials has evident advantages in identifying genotype-dependent responsiveness and adverse effects and has been advocated for despite such experimental designs remaining rare [15]. Most interestingly, however, GxD experiments have the potential to vastly improve the design of therapeutic trials, as the identification of disease-associated pathways altered in only a subset of genetic backgrounds can be used to predict that those genotypes, and those genotypes alone, will benefit from a drug targeting that pathway.

Here we follow a new experimental paradigm in which we identify GxD changes in gene expression and DNA methylation in a cohort of genetically diverse mouse strains, apply the pathway analysis from those studies to inform a literature-based selection of a predicted strain-specific drug intervention, and assess the molecular and phenotypic consequences of this intervention in a strain-specific manner (Fig 1). Specifically, we identified as a candidate genotype-specific therapy a Farnesoid X Receptor (FXR) agonist, GW4064 [16], based on the observation of strain-specific modulation of the FXR pathway in response to a high-fat high-carbohydrate “American” diet, and found that this drug not only has beneficial effects on the strain predicted to be responsive, but also shows strain-specific adverse responses in the strain predicted to be insensitive. In so doing, we have identified an example of both strain-specific disease phenotype mitigation and strain-specific deleterious side effects. This integrative approach thus opens the door to harnessing mouse genetics to improve preclinical assessment of both benefit and risk for genotype- and diet-specific drug candidates.

**Fig 1.**
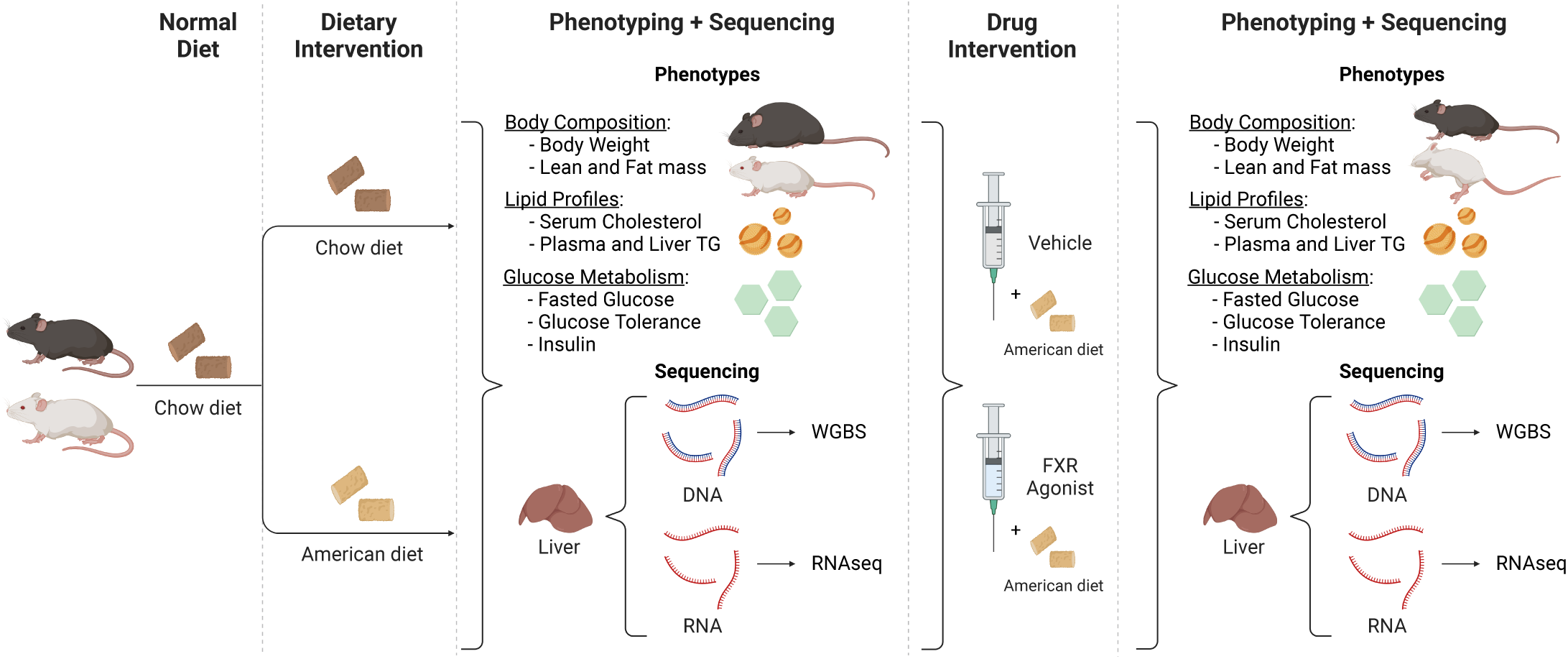
Schematic of experimental design. Genetically divergent mouse strains BL6, A/J, and NOD were exposed to either the American diet or a standard mouse chow. Phenotypic measurements were collected and WGBS and RNA-seq were performed on DNA and RNA from liver. Separately, BL6 and NOD mouse strains were exposed to an American diet and were treated with the FXR agonist GW4064 or control vehicle. Phenotypic measurements were collected and WGBS and RNA-seq were performed on DNA and RNA from liver. Comparison of differentially methylated and expressed genes between strains allows identification of common and strain-specific responses to the American diet and GW4064 treatment. Graphics created with BioRender.com.

## Results

### Characteristics of the mouse-diet experimental model

To elucidate how diverse genetic backgrounds differentially mediate phenotypic response to the environment, we designed epigenetic and transcriptomic analyses on three mouse genotypes exposed to two different diets. We selected three inbred founder strains from the Collaborative Cross project [17] and two diets with relevance to human nutrition: control laboratory chow (termed throughout as “standard”) and a high-fat, high-carbohydrate “American” diet designed to match typical Western nutritional intakes which have been found to be detrimental to metabolic health in both animal and human studies [7, 18]. Each of the three selected mouse strains has a different degree of phenotypic response to the American diet when compared to the standard diet [7]. BL6, the most commonly used laboratory strain and the source of the standard mouse reference genome, displays strong negative changes in metabolic phenotypes on the American diet, including large increases in body fat, hepatic triglycerides, and total cholesterol. These phenotypic changes are in line with previous literature showing that the BL6 strain was adversely affected by high-fat diets [19, 20]. In contrast, the A/J (AJ) and NOD/ShiLtJ (NOD) strains are more resistant to the American diet, with A/J mice displaying only mild changes in metabolic measurements and NOD showing moderate changes. This outcome is also in line with previous literature observations that the A/J strain was resistant to high-fat diets [21] and that NOD is a non-obese diabetic model [22].

The wide range of phenotypic response in our chosen combination of strains and diets provides a robust experimental framework for identifying epigenetic GxD interactions (Fig 1). First, differentially expressed genes (DE genes) that change with diet and differentially methylated regions (DMRs) denoting diet-dependent methylation changes can be identified for each strain individually. Next, by assessing the extent to which expression or methylation changes overlap between strains, "strain-specific" DE genes and DMRs can be identified that are altered on a genotype-by-diet basis.

### Strain-specific gene expression analysis reveals common and distinct targets and pathways

First, we performed RNA sequencing (RNA-seq) on liver tissue from of BL6, A/J, and NOD mice on standard and American diets (n=5 per group; n=4 per group for NOD). The liver was chosen for its relative tissue homogeneity as well as physiological relevance in metabolism and disease, including observed changes in hepatic triglyceride levels. We used kallisto to pseudo-align reads [23], utilizing personalized reference transcriptomes specific to each strain to minimize alignment bias from sequencing differences. Then, for each strain independently, we used edgeR/limma [24, 25] to identify genes that were DE (adjusted p-value < 0.05) between standard and American diet groups.

Our analysis identified 1,307 genes that are DE in at least one strain (Fig 2a and 2b, S1-3 Tables). Strikingly, nearly all DE genes are genotype-specific, with only 26 genes that are DE in all three strains (Fig 2b). BL6 has by far the most changes in expression (1,129 DE genes), while A/J displays minimal expression changes (75 DE genes), and NOD has an intermediate number of changes (245 DE genes). This result closely mirrors the degree of phenotypic diet response in these strains.

**Fig 2.**
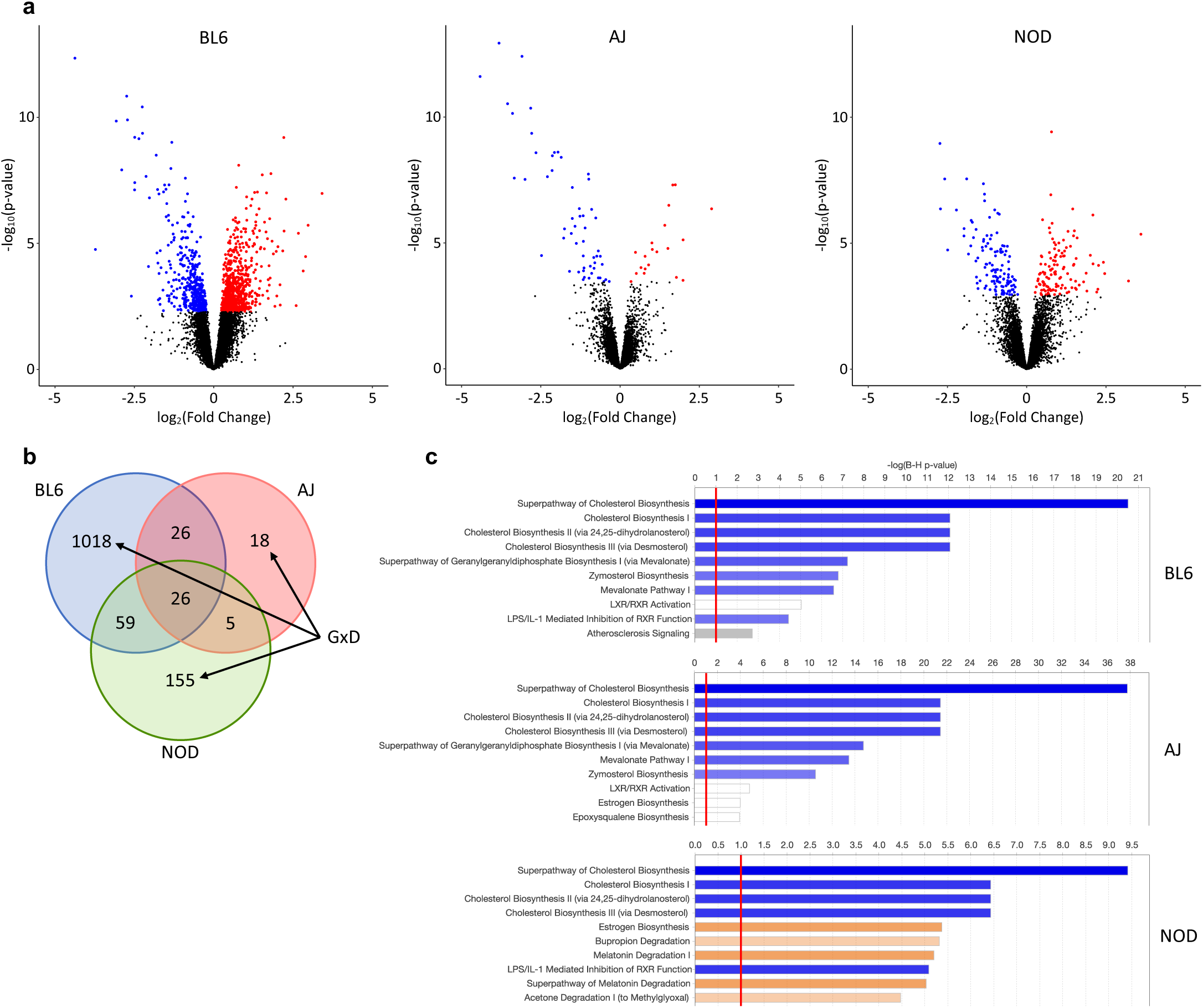
Gene-by-environment interactions of hepatic gene expression in BL6, A/J, and NOD mice. **a**, Volcano plots indicating significantly upregulated (red) and downregulated (blue) genes in BL6, A/J, and NOD, as determined by Benjamini-Hochberg adjusted p-value. **b**, Venn diagram of overlap between strains of differentially expressed genes (DEGs) upon diet exposure. Marked categories are strain-specific, highlighting potential gene-by-diet interactions. **c**, IPA results showing enriched pathways within diet DEGs for each strain; the top 10 most significantly enriched pathways with Benjamini-Hochberg FDR-adjusted p-values < 0.1 are shown.

To determine processes or pathways where gene expression is uniquely modulated by diet in each strain, we performed IPA on the DE genes identified in each strain. The top 10 most significantly enriched pathways (adjusted p-value < 0.1) for each strain are shown in Fig 2c and all significantly enriched pathways are shown in S1-3 Figs. Cholesterol biosynthesis pathways were highly enriched in all strains, followed by pathways unique to a subset of strains. For example, BL6 DE genes were uniquely enriched for atherosclerosis signaling, possibly reflecting the strain’s highly adverse phenotypic responses to the American diet. Interestingly, while A/J and NOD had relatively few DE genes compared to BL6, DE genes from these two strains were enriched for the estrogen biosynthesis pathway, which has been shown to play roles in regulating hepatic lipid metabolism [26] and is not significantly enriched in BL6 DE genes. Thus, these gene ontology results reveal potential epigenetic mechanisms by which genotype-specific regulation of gene expression confers protective or deleterious responses to the same diet in different strains.

### Applying GxD pathway analysis predicts strain-specific efficacy of metabolic drugs

The large GxD effects we observed in phenotypic and transcriptomic responses to the American diet indicate that genotype is a major factor in determining an individual’s sensitivity to environmental challenges. Notably, these findings significantly reinforced our hypothesis that an individual strain’s responsiveness to drug treatment, specifically one aimed at protecting against deleterious diet effects, could be equally genotype-dependent. Thus, we sought to determine if GxD pathways identified through our transcriptomic analysis as strain-specific could be utilized to predict certain drugs as having genotype-dependent efficacy.

To identify druggable gene targets which could elicit genotype-specific protection against diet-induced obesity-associated phenotypes in our mouse strains, we performed a regulatory network analysis utilizing IPA, further described in our methods section, to identify upstream transcriptional regulators which could explain observed changes in gene expression [27]. Of the master upstream regulators identified by this analysis, some were ubiquitously dysregulated by diet across all three strains, while others showed a large degree of strain-specificity (Fig 3a). We then searched for commercially available metabolic drugs with established efficacy and characterized mechanism of function in BL6, namely a protective effect against a high-fat diet, and asked whether any of them affected BL6-specific master upstream regulators.

**Fig 3.**
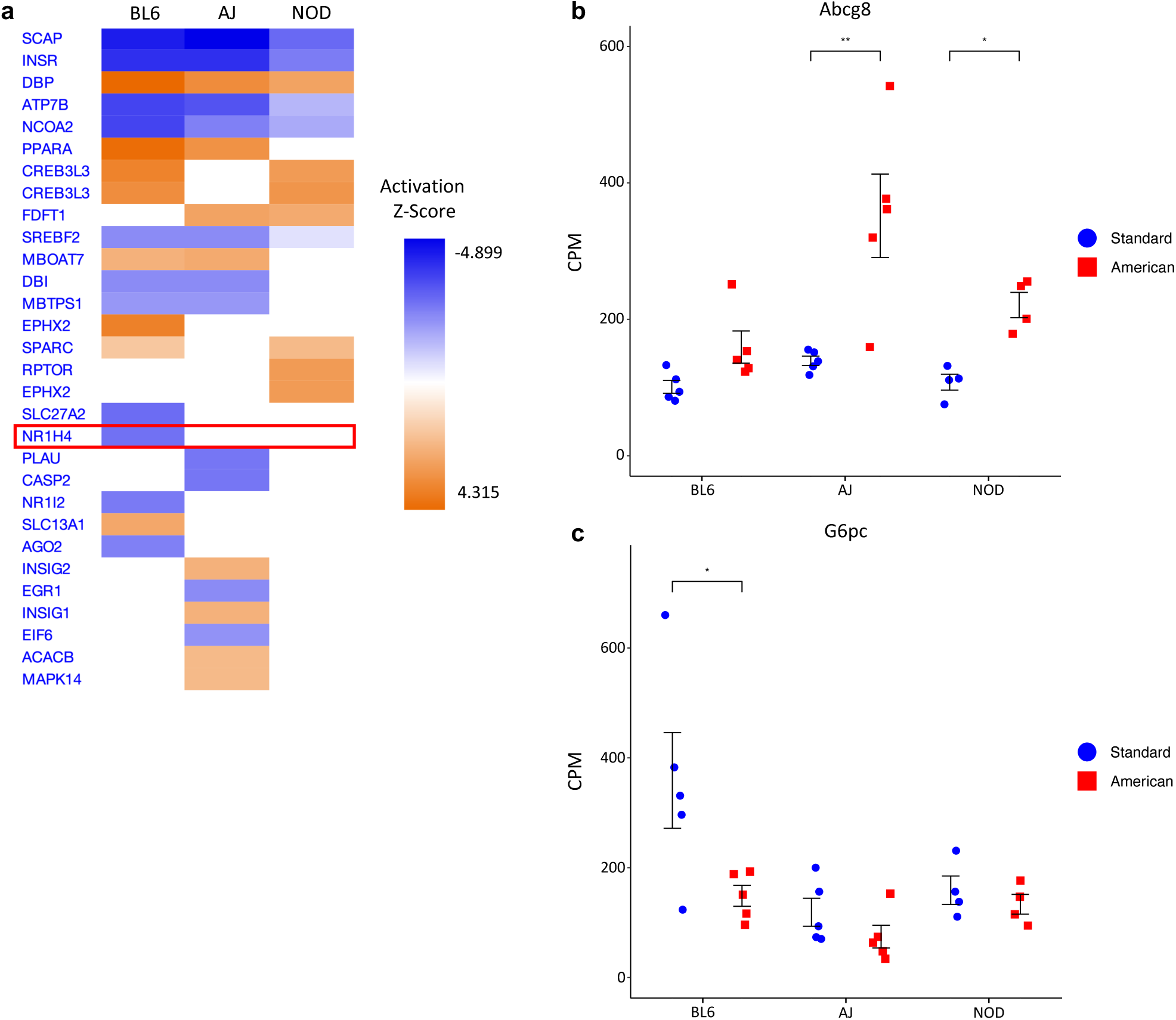
Regulatory network analysis identifies ubiquitous and strain-specific transcriptional regulators of diet response. **a**, Output of IPA regulatory network analysis identifying both common and strain-specific master upstream transcriptional regulators predicted based on diet DEGs in BL6, A/J, and NOD. The top 30 master upstream regulators ranked by z-score are shown. FXR (*Nr1h4*) is highlighted. **b**, Plot of *Abcg8* gene expression in BL6, A/J, and NOD mice on standard and American diets. **c**, Plot of *G6pc* gene expression in BL6, A/J, and NOD mice on standard and American diets. Error bars represent SE. * p < 0.05, ** p < 0.01, *** p < 0.001 from RNA-seq Benjamini-Hochberg FDR-adjusted p-values.

Based on these criteria, we selected GW4064, a well-documented FXR-agonist previously shown to prevent diet-induced obesity in BL6 mice [16]. FXR (*Nr1h4*) is a master upstream regulator predicted via our IPA regulatory network analysis to be downregulated in BL6 and unchanged in A/J and NOD (Fig 3a). As examples of individual genes affected by activation of FXR, GW4064 has been shown to upregulate *Abcg8*, a cholesterol transporter that is an indirect target of FXR [28]. In A/J and NOD mice, *Abcg8* expression was highly upregulated upon American diet exposure but was unchanged in BL6 (Fig 3b). Conversely, *G6pc*, another gene predicted by IPA to be activated by FXR, was downregulated upon American diet exposure in BL6, but not in A/J or NOD (Fig 3c). These results suggest that FXR pathway regulation is impaired in BL6 but not A/J or NOD mice. We thus reasoned that GW4064-induced activation of FXR might protect BL6 mice, but not NOD mice, from the deleterious effects of an American diet.

### FXR agonist GW4064 shows distinct GxD efficacy and toxicity

To test for genotype-specific responses to GW4064, we exposed a cohort of 15-week-old BL6 and NOD mice (n=5 per group) to the standard and American diet chow for 6 weeks. Mice on the American diet were treated with either GW4064 or vehicle via intraperitoneal injection; standard diet mice were treated with vehicle only. To characterize the health effects of diet and GW4064 treatment on these mice, we took a variety of metabolic measurements throughout the 6-week period, including lean and fat body mass, cholesterol levels, liver triglyceride levels, and glucose tolerance tests (S4 Table). Four NOD mice were euthanized early due to health issues during testing and were removed from subsequent analyses.

Strikingly, both American diet and GW4064 treatment exerted strongly strain-specific effects on several metabolic phenotypes (Fig 4a). BL6 mice exhibited a large increase in body fat on the American diet that GW4064 almost entirely reversed, consistent with previous reports and our earlier results (Fig 4a, left). In contrast, NOD mice did not exhibit any such increase in body fat, and thus had no phenotype to revert with GW4064 treatment. Even more notable were trends in cholesterol and liver triglycerides which were both increased in both strains upon exposure to the American diet but reverted upon GW4064 treatment only in BL6 mice (Fig 4a, center and right). That is, treatment of American-diet mice with GW4064 caused statistically significant decreases in cholesterol and liver triglycerides in BL6 alone, with little to no change observed in NOD mice. Hepatic histopathology also revealed strain-specific responses to GW4064 (S4 Fig, S5 Table). All mice from both strains on the American diet developed moderate to severe levels of hepatic lesions that were not seen in liver samples from mice from either strain when on the standard diet. This hepatic damage was prevented in all five BL6 mice on the American diet when also given GW4064. In contrast, one of three NOD mice on the American diet given GW4064 still exhibited hepatic damage. Observed hepatic damage mainly consisted of glycogenosis and lipidosis, which was low in all mice on the standard diet but increased significantly when mice were fed the American diet, irrespective of strain. Average levels of hepatic glycogenosis and lipidosis were significantly lower in both strains fed the American diet while also receiving GW4064, albeit levels remained slightly higher in NOD compared to BL6, especially for lipidosis which was consistent with hepatic triglyceride differences described earlier. Together, our results strongly indicate that the FXR agonist GW4064 has a genotype-specific effect on mice fed an American diet, namely beneficial responses in BL6 mice compared to minimal responses in NOD mice.

**Fig 4.**
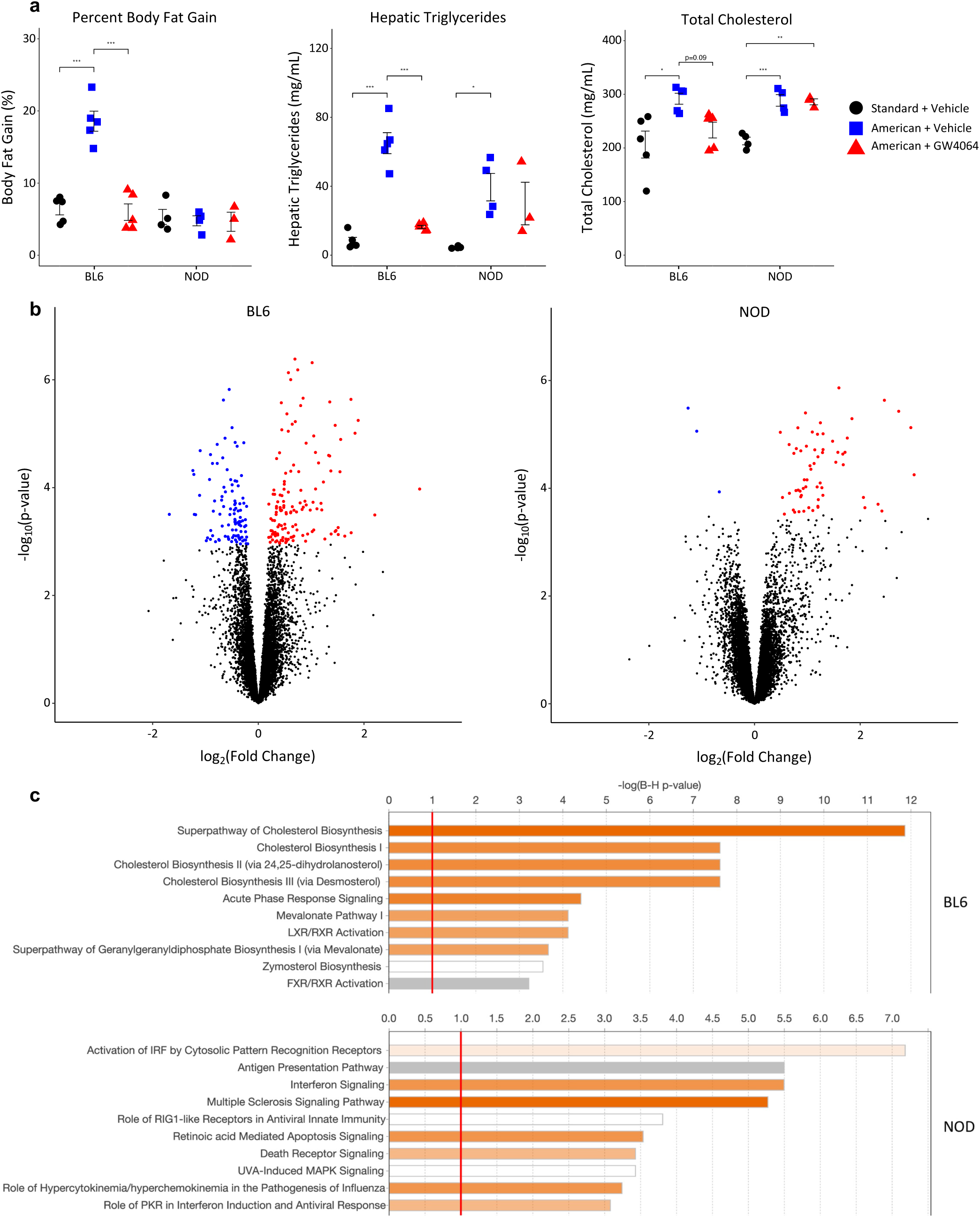
FXR agonist GW4064 elicits strain-specific responses in BL6 and NOD mice on the American diet. **a**, Phenotypic measurements of BL6 and NOD mice under three conditions tested (standard diet + vehicle, American diet + vehicle, American diet + GW4064). Phenotypes shown are percent body fat gain (left), hepatic triglyceride levels (middle), and total cholesterol (right). Error bars represent SE. * p < 0.05, ** p < 0.01, *** p < 0.001 from ANOVA between means within strains, with post-hoc Tukey HSD test. **b**, Volcano plots of liver gene expression indicating significantly upregulated (red) and downregulated (blue) genes in BL6 and NOD, as determined by Benjamini-Hochberg adjusted p-value. **c**, IPA results showing enriched pathways within drug DEGs for each strain; the top 10 most significantly enriched pathways with Benjamini-Hochberg FDR-adjusted p-values < 0.1 are shown.

To explore potential mechanisms underlying the genotype-specific response to GW4064, we performed RNA-seq on livers of BL6 and NOD mice on the American diet, under the vehicle or GW4064 treatment conditions, then identified DE genes between vehicle and drug for each strain (S6-7 Tables). BL6 mice had a larger number of DE genes compared to NOD, consistent with its stronger phenotypic response to GW4064 (Fig 4b). Furthermore, DE genes in BL6 and NOD are almost entirely different, with only 6 DE genes common to both strains compared to 231 and 58 which are specific to BL6 and NOD, respectively. This suggests highly disparate responses to GW4064 between these genotypes and mirrors our earlier observations of strongly genotype-specific responses to diet (Fig 2b). To elucidate whether unique biological processes are affected by GW4064 treatment in BL6 compared to NOD, we used IPA to examine gene ontology of DE genes. The top 10 most significantly enriched pathways (adjusted p-value < 0.1) for each strain are shown in Fig 4c and all significantly enriched pathways are shown in S5-6 Figs. BL6 DE genes are enriched in pathways highly relevant to metabolism, showing very strong enrichment in cholesterol biosynthesis in particular (Fig 4c and S5). Strikingly, NOD DE genes show no enrichment for these same metabolic pathways, but instead have strong enrichments for immune-related pathways including interferon signaling and activation of antiviral response pathways (Fig 4c and S6). This surprising and consistent enrichment in immune-related pathways within the NOD DE genes serves to potentially highlight that drug treatments, in addition to having genotype-specific benefits resulting from the return of dysregulated gene networks to normal function, may also have genotype-specific toxicities resulting from the augmentation of gene networks which were not dysregulated by the change in diet to begin with.

### Common and distinct targets and pathways in diet-associated DNA methylation changes

Given the capability of the epigenome, and in particular DNA methylation, to mediate phenotypic responses to both genetic and environmental factors, we were interested to see how methylation was altered in response to dietary changes in each of the three strains tested. To test this, we performed whole-genome bisulfite sequencing (WGBS) to assess DNA methylation in liver tissue of the same BL6, A/J, and NOD mouse individuals in which we performed RNA-seq (n=5 per group; n=3 per group for NOD). To accurately measure methylation in different mouse genotypes, which can be biased by alignment to incorrect reference genomes, we aligned WGBS reads to personalized reference genomes for each strain [29]. We then performed a permutation-based analysis for each strain to identify diet-associated candidate DMRs and further filtered these DMRs by removing those with less than a 10% mean methylation difference between the standard and American diets. Due to the exploratory nature of this analysis, we chose to further analyze all nominally significant DMRs which also pass the post-hoc methylation difference filtering. 1,316 DMRs were identified with a diet-associated methylation change in at least one strain (Fig 5a, S8-10 Tables). BL6 contains the highest number of diet DMRs (728), which may reflect its high phenotypic responsiveness to the American diet relative to the other two strains, while NOD exhibits an intermediate number of DMRs (367), and A/J has the fewest (299). As observed with diet-associated expression changes, this trend perfectly matches the degrees of phenotypic changes associated with each strain.

**Fig 5.**
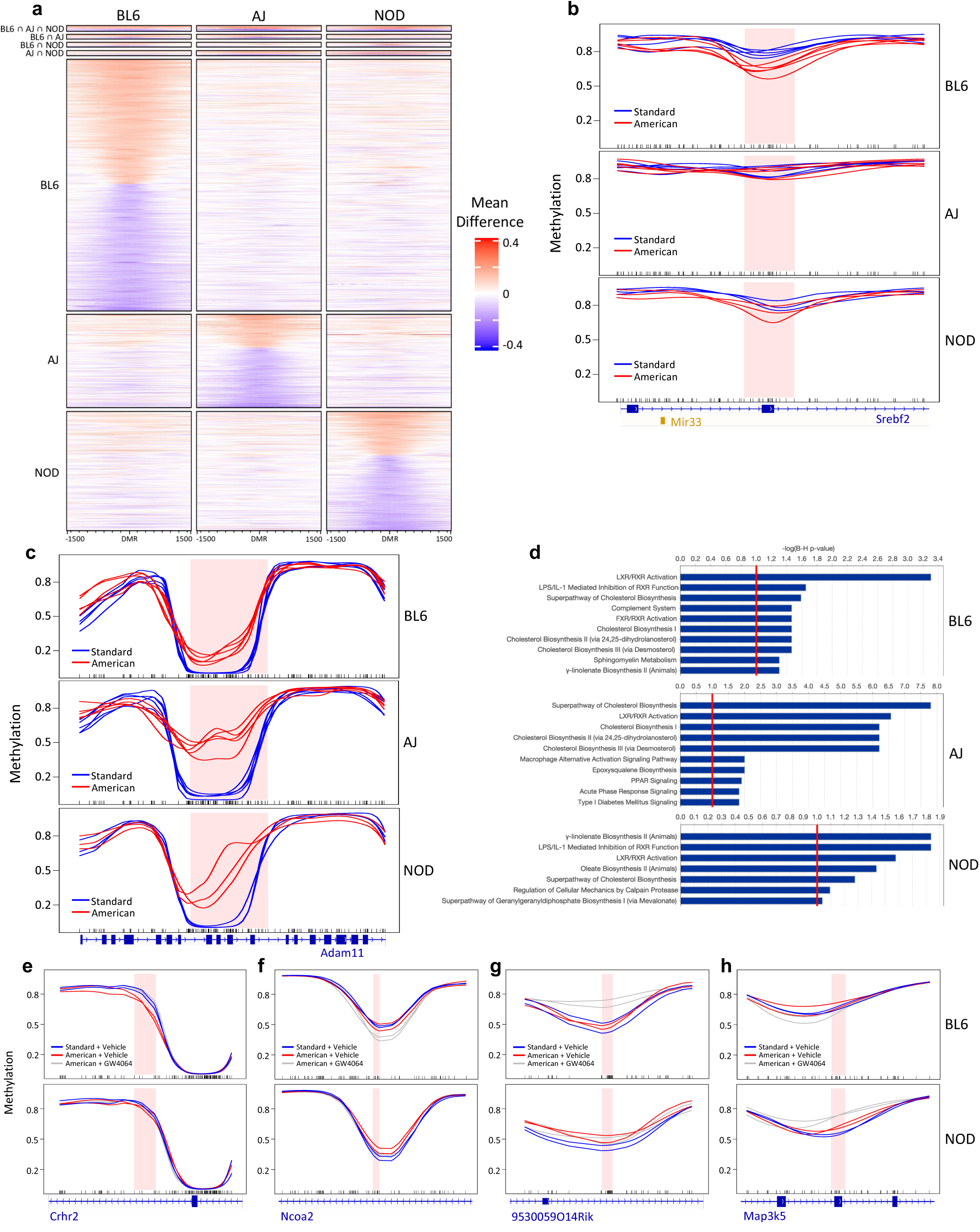
Gene-by-environment interactions in hepatic DNA methylation of BL6, A/J, and NOD mice. **a**, Heatmap of mean methylation differences between diets across a 3-kb window centered around 1,316 DMRs, grouped by strain uniqueness. DMRs in group 1 are nominally significant and have dietary mean difference > 10% in all three strains; groups 2 through 7 denote DMRs which are nominally significant and have mean difference > 10% in only one or two strains. **b**, Example of a strain-specific DMR, showing the BL6-specific hypomethylation of *Srebf2* and *Mir33* upon exposure to the American diet. **c**, Example of a strain-agnostic DMR, showing the ubiquitous hypermethylation of *Adam11* upon exposure to the American diet. **d**, IPA results showing enriched pathways within diet DMR-associated genes for each strain; the top 10 most enriched pathways with Benjamini-Hochberg FDR-adjusted p-values < 0.1 are shown. **e-h**, Examples of four distinct categories of diet- and drug-associated DMRs overlapping phenotypically relevant genes functionally implicated in various metabolic **(e-g)** or inflammatory **(h)** pathways.

Strikingly, of the 1,316 identified DMRs, only 17 regions (1.3%) show diet changes in the same direction in all three strains. The vast remainder of the genome is instead characterized by genotype-specific DMRs, with 1,258 (95.6%) showing diet change in only one strain (Fig 5a). One example of a genotype-specific DMR is a region in the *Srebf2* gene, which is involved in cholesterol homeostasis [30] and which becomes hypomethylated only in BL6 mice, but not A/J or NOD mice, upon American diet exposure (Fig 5b). As an example of a DMR that is common to all three strains, a region within the *Adam11* gene becomes hypermethylated in all strains on the American diet (Fig 5c). Notably, *Adam11* is also significantly overexpressed within the American diet of all three strains, potentially indicating a link between the methylation of this DMR and *Adam11* gene expression.

Using the ChIPSeeker R package, which performs annotation of genomic regions [31], we next examined the gene associations of DMRs. DMRs were associated with the promoter or gene body of 887 Ensembl genes; naturally, due to the genotype-specific nature of most DMRs, each strain was associated with a distinct set of genes, with only 19 DMR-associated genes common to all three strains. We used QIAGEN’s Ingenuity Pathway Analysis (IPA) to identify enriched pathways in DMR-associated genes for each strain (Fig 5d and S7-9). As we observed with diet-associated DE genes, cholesterol biosynthesis pathways and activation of the LXR/RXR pathway, which involves nuclear receptors with roles in lipid metabolism and transport [32, 33], were enriched in all three strains, while other pathways were unique to certain strains; for example, the superpathway of geranylgeranyldiphosphate biosynthesis, which has been implicated in high-fat diet-associated non-alcoholic fatty liver disease fibrosis [34], was only enriched in A/J and NOD, while BL6 was uniquely enriched for the complement system, which has been previously associated with obesity and may also play roles in the development of insulin resistance and diabetes mellitus [35]. Overall, our results indicate that, while certain dysregulated pathways appear to be universal or strain-agnostic effects of dietary intervention, there are many strongly genotype-specific methylation changes in distinct pathways that underlie the differential response of BL6, A/J, and NOD mouse strains to the same American-diet exposure.

### Genotype-specific methylation responses to diet-protective drug treatment

Finally, to examine methylation patterns that might contribute to the observed genotype-specific drug response, we performed WGBS on livers of the same BL6 and NOD mice from all three diet and drug treatment groups (standard diet with vehicle, American diet with vehicle, and American diet with GW4064). Using this exploratory small-sample WGBS study (n=2 mice per group), we sought to confirm the presence of phenotypically relevant methylation patterns over obesity- and metabolism-related genes. We first called 355 and 309 “drug DMRs” between vehicle- and GW4064-treated mice on the American diet for BL6 and NOD, respectively, which represent regions where GW4064 treatment elicits a significant methylation change in each strain (S11-12 Tables). Interestingly, of these DMRs, only 26 (7.32%) in BL6 and 15 (4.85%) in NOD also showed a diet-induced methylation change between standard and American diets (S13-14 Tables). This suggests that reversion of methylation changes caused by the American diet within BL6 account for only a subset of GW4064’s effects, and that, instead, GW4064 promotes many methylation changes at regions where BL6 would not otherwise respond to American diet exposure. These GW4064-dependent changes, in turn, could represent protective metabolic effects that BL6 mice cannot normally produce on their own.

We next examined NOD methylation over BL6 drug DMRs to determine how NOD mice might respond differently at these loci that undergo drug-induced changes in BL6. For 73 (20.6%) of the BL6 drug DMRs, BL6 but not NOD displays diet-induced methylation change which is then reverted with GW4064 treatment. For example, at the *Crhr2* gene, which plays roles in lipid and cholesterol metabolism and has been implicated in obesity [36, 37], BL6 has significant changes on the American diet which is restored to standard-diet levels with GW4064 treatment, while NOD has no change in any diet or treatment group (Fig 5e). Other patterns of genotype-specific methylation differences were also observed. For 44 (12.4%) of the BL6 drug DMRs, BL6 mice are hyper/hypomethylated relative to NOD mice on the standard diet, and this methylation does not change upon American diet exposure; GW4064 treatment induces a change in methylation to reach levels similar to those in NOD. This suggests that NOD’s intrinsic methylation level over these regions could be protective against the American diet. An example of this pattern is observed in the intron of *Ncoa2*, a gene with important roles in adipogenesis and lipid metabolism [38-40] (Fig 5f). At another DMR, over the ncRNA *9530059O14Rik*, NOD but not BL6 mice display hypermethylation upon American diet exposure (though the observed 8.8% methylation change is slightly under the 10% cutoff), while BL6 mice only gain methylation when treated with GW4064 (Fig 5g). This type of DMR could represent protective responses to the American diet in the NOD genotype that are not activated in BL6 mice, and that must instead be compensated for by GW4064 treatment. Interestingly, this DMR overlaps quantitative trait loci influencing obesity, hepatic cholesterol accumulation, and triglyceride concentrations [41, 42].

Lastly, we identified NOD drug DMRs over which no methylation changes are observed in BL6 nor between the American and standard diets in NOD mice. These regions could represent GW4064-induced methylation changes that contribute to the NOD-specific immune activation predicted from the pathway analysis of the previously described transcriptomic data. 288 (93.2%) of NOD drug DMRs fall within this category, and several can be functionally related back to the predicted drug-related immune response. As an example, a DMR following this pattern is observed over *Map3k5* (also known as *Ask1*), a gene which plays a role in apoptotic signaling and has been shown to be induced by inflammatory cytokines [43] (Fig 5h). Overall, these methylation and gene expression studies on GW4064-treated mice point towards a model in which genotype-specific methylation, including both intrinsic levels of methylation as well as disparate responses to an environmental exposure, can predetermine, in part, consequences of a diet or efficacy of a drug treatment for a given genotype.

## Discussion

In this study, we coupled diet exposure with a set of genetically diverse mouse strains to ascertain the effect of genetic variation on epigenetic and transcriptomic responses to the environment, as measured by DNA methylation and gene expression. While a small number of metabolic genes were commonly activated or repressed across all mouse strains, such as in cholesterol biosynthesis pathways, a far larger set of genes falling into unique metabolic regulatory networks were altered on a strain-unique basis. We also applied our observations on gene-environment interactions to test genotype-specific responses to the metabolic drug GW4064. We demonstrate that treatment with GW4064, while protective against the consequences of the high-fat American diet in BL6 mice, had limited to no effect on dysregulated metabolic phenotypes in NOD mice and may even induce NOD-specific toxicities. Together, these results suggest that any given diet or drug do not induce consistent pathways of response across a population; rather, each individual’s response is likely highly specific to that individual and is governed by the complex interactions between that individual’s unique set of genetic variants.

FXR activation is not the only potential candidate genotype-specific drug which has been identified through this analysis. Inhibitors which target *Slc13a1* and agonists which target *Nr1i2* or *Ago2* are predicted to have a similar BL6-specific protective effect against diet-induced obesity. Additionally, inhibitors of *Insig1*, *Insig2*, *Acacb*, or *Mapk14* and agonists of *Plau*, *Casp2*, *Egr1*, or *Eif6* are predicted to confer an A/J-specific protective effect. Finally, inhibitors of *Rptor* could have a NOD-specific protective effect. Perhaps most importantly, our integrative analysis also identifies pathways that may be druggable to confer a genotype-agnostic protective effect across all three tested strains. These include inhibitors which target *Dbp* and agonists which target *Scap*, *Insr*, *Atp7b*, or *Ncoa2*.

It should be noted that “strain-specific” is a term necessary only for our pilot study, which is limited to three strains. With more genotypes – for example, Collaborative Cross mice where alleles can be mapped with high resolution [44] – strain-unique DMRs resolve into “non-conserved” or “genotype-specific” DMRs present in a proportion of strains, which can be mapped to a genetic variant. In addition, these DMRs can, in a sense, be treated as a more granular substitute for physiological phenotypes, which are few and broadly controlled. In this case it is plausible that one large mapping panel could resolve a multitude of high-resolution genome-epigenome effects at once. This is an exciting prospect for any field, including dietetics, which would otherwise require the analysis of many complex interactions across heavily intertwined gene networks. Hypothetically, diets could even be reduced to their individual components, allowing for even more detailed association of genotype with one nutrient of interest.

In addition, this pilot study is limited in sample size which subsequently reduces statistical power, particularly for the identification of diet- and drug-associated DE genes and DMRs. Genome-wide corrections were applied when identifying DE genes, though only nominal p-values were utilized during DMR finding in order to broaden results. Future studies attempting to utilize a similar experimental and analytical methodology to identify transcriptomic and epigenetic drivers of genotype-specific phenotypic responses to environmental perturbations should seek to have additional biological replicates to overcome the stringency of these genome-wide corrections.

GW4064 has not been extensively explored in a clinical setting, largely due to its limited bioavailability and concerns that the presence of a stilbene group may cause hepatic toxicity as demonstrated previously in rats [45, 46]. In the present study, GW4064 induced hepatic damage and led to abnormal upregulation of immune and inflammatory response genes only in NOD mice, while BL6 mice demonstrated no adverse effects. These results have implications for how drug screening and preclinical animal trials could be performed. Specifically, they suggest that testing a diverse panel of mouse strains will prove far more valuable for identifying both genotype-specific response and genotype-specific adverse effects compared to common experimental designs where compounds are tested on a single strain of laboratory mouse. Such an approach could identify drug candidates that, while appearing ineffective across the general population, have high efficacy for a subset of individuals, and can explain genetic risk factors underlying rare adverse events. We propose that utilizing such experimental designs will become the paradigm going forward, as we expect that doing so will greatly expand the pool of gene targets for clinical study.

## Methods

### Sample and animal information

For the diet study, 4-week-old male A/J, BL6, and NOD mice were obtained from The Jackson Laboratory (Bar Harbor, ME) and acclimated for 2 weeks on a standard laboratory diet, then randomly assigned to standard or American diet groups, with five mice per strain and diet. Mice were fed their assigned diet in powdered form for roughly 6 months (24 weeks). Mice were housed at the University of North Carolina during the first 4 months for analysis of body composition, metabolic rate, and physical activity. Mice were transferred to North Carolina State University for the final 2 months of diet exposure, where they also underwent glucose tolerance testing (GTT), necropsy, and tissue collection, as described in [7]. Mice were housed five per cage and maintained at 22°C under a 12-hr light cycle; they were maintained, and protocols were followed in accordance with the University of North Carolina, North Carolina State University, and Texas A&M University Institution Animal Care and Use Committee guidelines. Mice were killed with carbon dioxide, and tissues were flash frozen in liquid nitrogen or fixed in formalin. For the GW4064 drug study, 9-week-old male BL6 and NOD mice were obtained from The Jackson Laboratory and acclimated for 6 weeks on a standard laboratory diet, then randomly assigned to standard-vehicle, American-vehicle, or American-GW4064 treatment groups, with five mice per strain and treatment. Four NOD mice, one on standard diet, one on American diet with vehicle, and 2 on American diet with GW4064, were euthanized early due health issues during testing. Mice were fed their assigned diet in pelleted form for roughly 6 weeks and treated with vehicle or GW4064 as described below (see “GW4064 formulation and treatment” section). Mice were housed at Texas A&M University at five per cage and maintained at 22°C under a 12-hr light cycle; protocols were followed in accordance with Texas A&M University Institution Animal Care and Use Committee guidelines. Mice were killed with carbon dioxide, and tissues were flash frozen in liquid nitrogen or fixed in formalin.

### Diet composition

Powdered and pelleted diets were designed in collaboration with Research Diets (New Brunswick, NJ); the American diet (D12052705) was based on the US Department of Agriculture’s 2008 Dietary Assessment of Major Food Trends, as described in [7]. A purified control mouse diet (D12052701, powder; D17031601, pellet; Research Diets) was used as a standard diet for comparison to the American diet. Diets were designed to recapitulate human diets as closely as possible, matching macronutrient ratio, fiber content, types of ingredients, and fatty acid ratios to the human diets. Accordingly, nutrient sources were selected to match intakes of human diets, e.g. beef protein to match red meat intake in the American diet.

### GW4064 formulation and treatment

GW4064 was purchased from Cayman Chemical Company (Catalog Number 10006611) and stored at -20°C in 20mg aliquots. Preparation of GW4064 solution for animal administration was performed on the day of injection. 20mg of GW4064 was first dissolved in 1000µL of 99.5% DMSO, then diluted with 1985µL of water to reduce DMSO concentration. 16µL of TWEEN 80 was then added to return GW4064 to solution. Vehicle solution was prepared with DMSO, water, and TWEEN 80 without GW4064. 50mg/kg of GW4064 solution, and equivalent volumes of vehicle, were administered to mice via intraperitoneal injection twice a week.

### DNA extraction and whole-genome bisulfite sequencing

Genomic DNA was extracted from liver samples using the Qiagen DNEasy kit, with an additional RNase incubation step (50 µg/sample, 30 minutes) prior to column application to remove RNA. For the American vs standard diet comparison, whole genome bisulfite sequencing single indexed libraries were generated using NEBNext Ultra DNA library Prep kit for Illumina (New England BioLabs) according to the manufacturer’s instructions with modifications. 500ng input gDNA was quantified by Qubit dsDNA BR assay (Invitrogen) and spiked with 1% unmethylated Lambda DNA (Promega, cat # D1521) to monitor bisulfite conversion efficiency. Input gDNA was fragmented by Covaris S220 or LE220 Focused-ultrasonicator to an average insert size of 350bp. Size selection to isolate insert sizes of 300-400bp was performed using AMPure XP beads. The EZ DNA Methylation-Gold Kit or EZ DNA Methylation-Lightning Kit (Zymo cat#D5005, cat#D5030) were used to bisulfite convert samples after size selection following the manufacturer’s instructions. Amplification was performed after the bisulfite conversion using Kapa Hifi Uracil+ (Kapa Biosystems, cat# KK282) polymerase using the following cycling conditions: 98°C 45s / 8cycles: 98°C 15s, 65°C 30s, 72°C 30s / 72°C 1 min. AMPure cleaned-up libraries were run on 2100 Bioanalyzer (Agilent) High-Sensitivity DNA assay, samples were also run on Bioanalyzer after shearing and size selection for quality control purposes. Libraries were quantified by qPCR using the Library Quantification Kit for Illumina sequencing platforms (KAPA Biosystems, cat#KK4824) using 7900HT Real-Time PCR System (Applied Biosystems). WGBS libraries were sequenced on an Illumina HiSeq2000 or HiSeq2500 instrument using 100bp paired-end indexed reads (v3 chemistry, BL6 and A/J samples) or 125bp paired-end indexed reads (v4 chemistry, NOD samples) with 10% PhiX spike-in. For the GW4064 drug study, the above protocol was followed with the following modifications: libraries were dual indexed, size selection was performed using SPRIselect beads, qPCR quantification was performed using the CFX384 Real-time system (BioRad), WGBS libraries were sequenced on an Illumina NovaSeq6000 instrument using 150bp paired-end dual indexed reads (S4 flowcell, version 1.5 reagents) with 5% PhiX spike-in.

### WGBS read alignment

TrimGalore (v0.6.6) was used to perform adapter removal and quality trimming of sequencing reads. In order to accurately estimate methylation while accounting for strain differences in genomic sequence, samples from BL6, A/J, and NOD were aligned to their respective reference genomes, obtained from UNC Systems Genetics (build 37), as we have previously described [29]. Alignment was performed using Bismark (v0.23.0) and Bowtie2 (v2.9.2). Reference genomes were combined with the λ phage genome for measurement of conversion efficiency. The Bismark function deduplicate_bismark was then used to remove PCR duplicates. M-bias plots were generated using bismark_methylation_extractor with the --mbias_only flag in order to identify the positions of biased CpG sites most commonly resulting from library prep end-repair. bismark_methylation_extractor was then used to extract methylation values. For BL6 and A/J samples, 8 and 3 bp from the 5’ and 3’ ends of read 1, respectively, and 12 and 5 bp from the 5’ and 3’ ends of read 2 were removed using the --ignore, --ignore_3prime, --ignore_r2, and --ignore_3prime_r2 flags based on M-bias results. For NOD samples, 5 and 3 bp from the 5’ and 3’ ends of read 1 and 15 and 5 bp from the 5’ and 3’ ends of read 2 were removed. CpG positions from A/J and NOD were converted into the BL6 (mm9) reference coordinate system using modmap [47]. These methylation values were used as input in the subsequent differential methylation analysis.

### Differential methylation analysis

Raw CpG methylation data from the cytosine reports output by bismark_methylation_extractor were imported into R version 3.6.1 using the read.bismark function of bsseq (v1.22.0) [48]. DMR identification was performed using dmrseq (v1.6.0) [49] with the default DMR-finding parameters except for the following: minNumRegion = 3, maxPerms = 100. For diet DMRs, the raw BSmooth object was subset to CpGs where coverage was greater than 2x in 4 out of 5 samples from each strain/diet, except for NOD samples in which coverage had to be greater than 2x in 2 out of the 3 samples from each diet. dmrseq was run using diet as the test covariate separately for each strain. Significant DMRs were defined as those with a dmrseq nominal p-value < 0.05 and a smoothed methylation difference between the groups, averaged across the entire DMR, of greater than 10%. To perform this calculation, CpG data were smoothed using the BSmooth/bsseq package (v1.22.0) with the default DMR-finding parameters (ns = 70, h = 1,000, maxGap = 10e8). Smoothing was performed over common and strain-unique CpGs to allow comparison of imputed methylation values across such sites, and then subset by coverage as previously described. To identify DMRs within the drug study, the same import, testing, and significance procedures and parameters as employed for the diet study were utilized, with a few differences. The raw BSmooth object was subset to CpGs where coverage was greater than 2x in all 12 samples. Two separate tests were run using dmrseq for each strain, one between the standard and American diet both given the vehicle and one between the American diet given the vehicle and the American diet given GW4064. Note that one mouse which was euthanized early was included in the drug DMR analysis (NOD sample on the American diet with GW4064 treatment).

### DMR annotation

DMR annotation was performed with ChIPSeeker version 1.22.1 [31]. Regions were annotated using the mm9 transcript database (TxDb.Mmusculus.UCSC.mm9.knownGene R package version 3.2.2) and the genome wide annotation for mouse (org.Mm.eg.db R package version 3.8.2) with the promoter region defined as 3-kb upstream or downstream of the transcription start site. DMR-associated genes for each strain were defined as genes annotated for a non-intergenic DMR with at least 10% standard-American methylation difference in that strain. CpG islands were obtained via the UCSC Genome Browser. CpG shores were defined as the 2-kb regions upstream and downstream of a CpG island.

### RNA extraction and RNA-seq

RNA was isolated from mouse liver using a Maxwell 16 LEV simplyRNA kit (Promega). For the American vs standard diet comparison strand-specific mRNA libraries were generated using the TruSeq Stranded mRNA protocol (Illumina, cat# RS-122-2101). Libraries were performed following the manufacturer’s protocol (Illumina, Part#15031050) with minor modifications. The input was 500ng (BL6 and NOD samples) or 2µg (A/J samples) and samples were fragmented for 6 min. The following PCR cycling conditions were used: 98°C 30s / 13 (BL6 and NOD samples) or 12 (A/J samples) cycles: 98°C 10s, 60°C 30s, 72°C 30s / 72°C 5 min. Stranded mRNA libraries were sequenced on an Illumina HiSeq4000 (BL6 and NOD samples) or HiSeq2500 (v4 chemistry; A/J samples) instrument using 75bp (BL6 and NOD samples) or 70bp (A/J samples) paired-end indexed reads and 1% of PhiX control. For the GW4064 drug study, strand-specific mRNA libraries were generated using the NEBNext Ultra II Directional RNA library prep Kit for Illumina (New England BioLabs #E7760), and mRNA was isolated using Poly(A) mRNA magnetic isolation module (New England BioLabs #E7490). The preparation of libraries followed the manufacturer’s protocol (Version 2.2 05/19). The input was 500ng and samples were fragmented for 15 min for RNA insert size of ∼200 bp. The following PCR cycling conditions were used: 98°C 30s / 8 cycles: 98°C 10s, 65°C 75s / 65°C 5 min. Stranded mRNA libraries were sequenced on an Illumina Hiseq4000 instrument using 48bp paired-end dual-indexed reads and 1% PhiX control.

### RNA-seq read alignment, quantification, and analysis

RNA-seq reads were quantified using the kallisto program (v0.46.1) [23], which uses pseudoalignment to match reads with target genes. cDNA FASTA files for BL6, A/J, and NOD genomes were obtained from The Jackson Laboratory. Each file was used to generate a strain-specific reference index to which reads from the corresponding strain’s samples were pseudoaligned and gene abundances estimated. RNA-seq data were analyzed in R version 3.6 using the edgeR (v3.28.1) and limma (v3.42.2) packages [24, 25]. Gene and transcript IDs were obtained from Ensembl release 101 and used as a target-mapping key to summarize kallisto abundance data at the gene level. Genes were filtered to those with a CPM (counts per million) greater than 1 in all 28 samples for the diet comparison and all 26 samples for the drug comparison; note that all mice which were euthanized early were removed prior to analysis. Normalization factors for library sizes were calculated with edgeR. A contrast matrix was designed to look for differential expression between standard and American diets in each strain individually. Raw counts were transformed to log-CPM values using the voom function from limma, then linear modelling was performed according to the contrast matrix to identify differentially expressed genes. Differentially expressed genes were defined as those with Benjamini-Hochberg FDR-adjusted p-values less than 0.05.

### Ingenuity Pathway Analysis – Enrichment Analyses

QIAGEN’s Ingenuity Pathway Analysis software was utilized to perform pathway enrichment analyses of RNA-seq differential gene expression and WGBS differential methylation data [27]. This enrichment analysis was performed on eight separate gene sets originating from the following three analyses: (1) differentially methylated genes associated with the change from standard mouse chow to the American diet from each of BL6, A/J, and NOD strains (three gene sets); (2) differentially expressed genes associated with the change from standard mouse chow to the American diet from each of BL6, A/J, and NOD strains (three gene sets); (3) differentially expressed genes associated with GW4064 treatment while on the American diet from the BL6 and NOD strains (two gene sets). For the gene sets of diet-induced differential methylation, expression analyses were performed using all genes associated with significant DMRs via ChIPSeeker. Default analysis settings were used except for the following: species was set to mouse only and tissues and cell lines were set to hepatocytes and liver. For the gene sets of diet-induced differential expression, an expression analysis was performed using the Log2FC values and the FDR-adjusted p-values as inputs. Cutoffs of Log2FC >= 1 and q < 0.05 were used. Default analysis settings were used except for the following: species was set to mouse only and tissues and cell lines were set to hepatocytes and liver. For the genes sets of drug-induced differential expression, an expression analysis was performed using the Log2FC values and the FDR-adjusted p-values as inputs. Cutoffs of Log2FC >= 0.5 and q < 0.05 were used. Default analysis settings were used except for the following: species was set to mouse only and tissues and cell lines were set to hepatocytes and liver.

### Ingenuity Pathway Analysis – Causal Network Analysis

QIAGEN’s Ingenuity Pathway Analysis software was also utilized to perform Causal Network Analysis on differentially expressed genes associated with the change from standard mouse chow to the American diet from each of BL6, A/J, and NOD strains. Separately, an expression analysis with the settings and inputs described above were performed on the gene sets from each of these strains. A comparison analysis was then run between the three expression analyses. Information from the Causal Networks section of the Upstream Analysis tab is presented in this paper.

### Liver pathology

Liver samples were fresh fixed in 10% neutral buffered formalin before being processed in a Leica ASP300 tissue processor for paraffin embedding by the Texas A&M Rodent Preclinical Phenotyping Core. After embedding, 5µm sections were cut on a Leica 2165 rotary microtome and sections H&E stained on a Leica HistoCore SPECTRA ST Stainer. After cover-slipping, slides were scored blinded by a board-certified veterinary pathologist. Severity of glycogenosis and lipidosis was scored on a 0-4 scale with 0 = normal and 4 = severe. Note that none of the four mice which were euthanized early were included in this analysis.

## Acknowledgements

This work was funded by NIH grants RM1HG008529 (to A.P.F. and D.W.T.), DP1DK119129 (to A.P.F. and D.W.T.)

## Supplementary Information

**S1 Fig. Pathway enrichments of BL6 diet DEGs.** IPA results showing enriched pathways within diet DEGs for BL6. All significantly enriched pathways with Benjamini-Hochberg FDR-adjusted p-values < 0.1 are shown.

**S2 Fig. Pathway enrichments of A/J diet DEGs.** IPA results showing enriched pathways within diet DEGs for A/J. All significantly enriched pathways with Benjamini-Hochberg FDR-adjusted p-values < 0.1 are shown.

**S3 Fig. Pathway enrichments of NOD diet DEGs.** IPA results showing enriched pathways within diet DEGs for NOD. All significantly enriched pathways with Benjamini-Hochberg FDR-adjusted p-values < 0.1 are shown.

**S4 Fig. Histological analysis of liver samples. a**, Scores for glycogenosis and lipidosis in each diet/treatment group. **b-g**, Representative liver histology images for: **b**, BL6 on standard diet with vehicle; **c**, BL6 on American diet with vehicle; **d**, BL6 on American diet with GW4064 treatment; **e**, NOD on standard diet with vehicle; **f**, NOD on American diet with vehicle; and **g**, NOD on American diet with GW4064 treatment. Bars in **b-g** are 200µm.

**S5 Fig. Pathway enrichments of BL6 drug DEGs.** IPA results showing enriched pathways within drug DEGs for BL6. All significantly enriched pathways with Benjamini-Hochberg FDR-adjusted p-values < 0.1 are shown.

**S6 Fig. Pathway enrichments of NOD drug DEGs.** IPA results showing enriched pathways within drug DEGs for NOD. All significantly enriched pathways with Benjamini-Hochberg FDR-adjusted p-values < 0.1 are shown.

**S7 Fig. Pathway enrichments of BL6 diet DMRs.** IPA results showing enriched pathways within diet DMRs for BL6. All significantly enriched pathways with Benjamini-Hochberg FDR-adjusted p-values < 0.1 are shown.

**S8 Fig. Pathway enrichments of A/J diet DMRs.** IPA results showing enriched pathways within diet DMRs for A/J. All significantly enriched pathways with Benjamini-Hochberg FDR-adjusted p-values < 0.1 are shown.

**S9 Fig. Pathway enrichments of NOD diet DMRs.** IPA results showing enriched pathways within diet DMRs for NOD. All significantly enriched pathways with Benjamini-Hochberg FDR-adjusted p-values < 0.1 are shown.

**S1 Table. BL6 diet DEGs.** Differentially expressed genes between the American and standard diet of BL6 mice.

**S2 Table. A/J diet DEGs.** Differentially expressed genes between the American and standard diet of A/J mice.

**S3 Table. NOD diet DEGs.** Differentially expressed genes between the American and standard diet of NOD mice.

**S4 Table. Drug treatment phenotypes.** Mouse phenotype information for BL6 and NOD mice with GW4064 + American diet and vehicle + standard diet exposures.

**S5 Table. Drug treatment hepatic histopathology.** Hepatic histopathology severity scoring of BL6 and NOD mice with GW4064 + American diet and vehicle + standard diet exposures.

**S6 Table. BL6 drug DEGs.** Differentially expressed genes between GW4064 + American diet and vehicle + American diet of BL6 mice.

**S7 Table. NOD drug DEGs.** Differentially expressed genes between GW4064 + American diet and vehicle + American diet of NOD mice.

**S8 Table. BL6 diet DMRs.** Differentially methylated regions between the American and standard diet of BL6 mice.

**S9 Table. A/J diet DMRs.** Differentially methylated regions between the American and standard diet of A/J mice.

**S10 Table. NOD diet DMRs.** Differentially methylated regions between the American and standard diet of NOD mice.

**S11 Table. BL6 drug DMRs.** Differentially methylated regions between the GW4064 + American diet and vehicle + standard diet of BL6 mice.

**S12 Table. NOD drug DMRs.** Differentially methylated regions between the GW4064 + American diet and vehicle + standard diet of NOD mice.

**S13 Table. BL6 diet DMRs with vehicle.** Differentially methylated regions between the vehicle + American diet and vehicle + standard diet of BL6 mice.

**S14 Table. NOD diet DMRs with vehicle.** Differentially methylated regions between the vehicle + American diet and vehicle + standard diet of NOD mice.

